# The Effect of Horizontal Gene Transfer on the Dynamics of Antibiotic Drug Resistance in a Unicellular Population with a Dynamic Fitness Landscape, Repression and De-repression

**DOI:** 10.1101/022012

**Authors:** Yoav Atsmon-Raz, Nathaniel Wagner, Emanuel David Tannenbaum

## Abstract

Antibiotic drug resistance spreads through horizontal gene transfer (HGT) via bacterial conjugation in unicellular populations of bacteria. Consequently, the efficiency of antibiotics is limited and the expected “grace period” of novel antibiotics is typically quite short. One of the mechanisms that allow the accelerated adaptation of bacteria to antibiotics is bacterial conjugation. However, bacterial conjugation is regulated by several biological factors, with one of the most important ones being repression and derepression.

In recent work, we have studied the effects that repression and de-repression on the mutation-selection balance of an HGT-enabled bacterial population in a static environment. Two of our main findings were that conjugation has a deleterious effect on the mean fitness of the population and that repression is expected to allow a restoration of the fitness cost due to plasmid hosting.

Here, we consider the effect that conjugation-mediated HGT has on the speed of adaptation in a dynamic environment and the effect that repression will have on the dynamics of antibiotic drug resistance. We find that, the effect of repression is dynamic in its possible outcome, that a conjugators to non-conjugators phase transition exists in a dynamic landscape as we have previously found for a static landscape and we quantify the time required for a unicellular population to adapt to a new antibiotic in a periodically changing fitness landscape. Our results also confirmed that HGT accelerates adaptation for a population of prokaryotes which agrees with current knowledge, that HGT rates increase when a population is put under stress.

## INTRODUCTION

Horizontal Gene Transfer (HGT) is defined as the transfer or exchange of genetic information between two organisms, that does not involve the transmission of genetic information from parent to daughter as a result of replication (Ochman *et al.* 2000; Gogarten and Townsend 2005; French 2010b). Since its discovery in the first half of the last century (Tatum and Lederberg 1947) in *Escherichia Coli* it was observed in Archaea (Zhaxybayeva and Doolittle 2011), Prokaryotes (Ochman *et al.* 2000), and Eukaryotes (Kidwell 1993; Currat and Excoffier 2011). Furthermore, it has been discovered that HGT can even occur among the three domains of biology (Zhaxybayeva and Doolittle 2011; Syvanen 2012; Bruto *et al.* 2013).

Consequently, HGT has become a subject of great interest for molecular and evolutionary biologists, due to accumulating evidence that suggests that HGT plays a large role in the re-shaping of prokaryotic genomes (Hiramatsu 1998; Ochman *et al.* 2000; Cohen *et al.* 2005) and for being the main reason for the rapid outspread of antibiotic drug resistance in bacterial populations (De La Cruz and Davies 2000; Ochman *et al.* 2000; Walsh 2000; Weldhagen 2004; Cohen *et al.* 2005; Hanssen and Ericson Sollid 2006; Robicsek *et al.* 2006; Roberts 2008). The implications that arise from the latter of these reasons have brought the realization that HGT may have severe consequences for public health care worldwide due to the rapid outspread of resistance genes between different species of bacteria (Palumbi 2001).

Prokaryotic genomes typically consist of a single, large, circular chromosome, in addition to smaller, circular chromosomes known as *plasmids*. Bacterial plasmids can move between bacteria, often between bacteria of different strains, via a process known as *conjugation* (Cruz and Lanka 1998; Grillot-Courvalin *et al.* 1998; S. Russi 2008; Carattoli 2013). When these plasmids contain genes for an adaptive trait such as antibiotic drug resistance (Van Meervenne *et al.* 2012; Clewell 2014), translation of Colicins (Cascales *et al.* 2007) or resistance to heavy metals (Mergeay *et al.* 1985; Silver and Misra 1988), then conjugation can significantly speed the outspread of that trait throughout a population (Novozhilov *et al.* 2005; Mc Ginty *et al.* 2013; Schulte *et al.* 2013). Therefore, conjugation is considered to be one of the most important forms of HGT and has a major role in the emergence of antibiotic drug resistance.

The most well studied bacterial conjugation system is the F^+^/F^−^ system (S. Russi 2008) in the *Escherichia Coli* K-12 strain. It also is very well known for its ability to increase the outspread of antibiotic drug resistance by the integration of relevant genes into the plasmid’s genome (Van Meervenne *et al.* 2012; Clewell 2014). In the F^+^/F^−^ system, a bacterium containing an F-plasmid fuses with a bacterium lacking the F-plasmid. The bacterium containing the F-plasmid is denoted as an F^+^ bacterium, while the bacterium that does not contain this plasmid is denoted as an F^−^ bacterium. When an F^+^ bacterium meets an F^−^ bacterium, it transfers one of the strands of the F-plasmid to the F^−^ bacterium via a pilus. Once a strand of the F-plasmid has been transferred from the F^+^ bacterium to the F^−^ bacterium, a copy of the plasmid in both cells is produced by daughter strand synthesis using the DNA template strands. The F^−^ bacterium then becomes an F^+^ bacterium that transcribes its own pilus, and is able to transfer the F^−^plasmid to other bacteria in the population (Figure 1).

**Fig 1.**
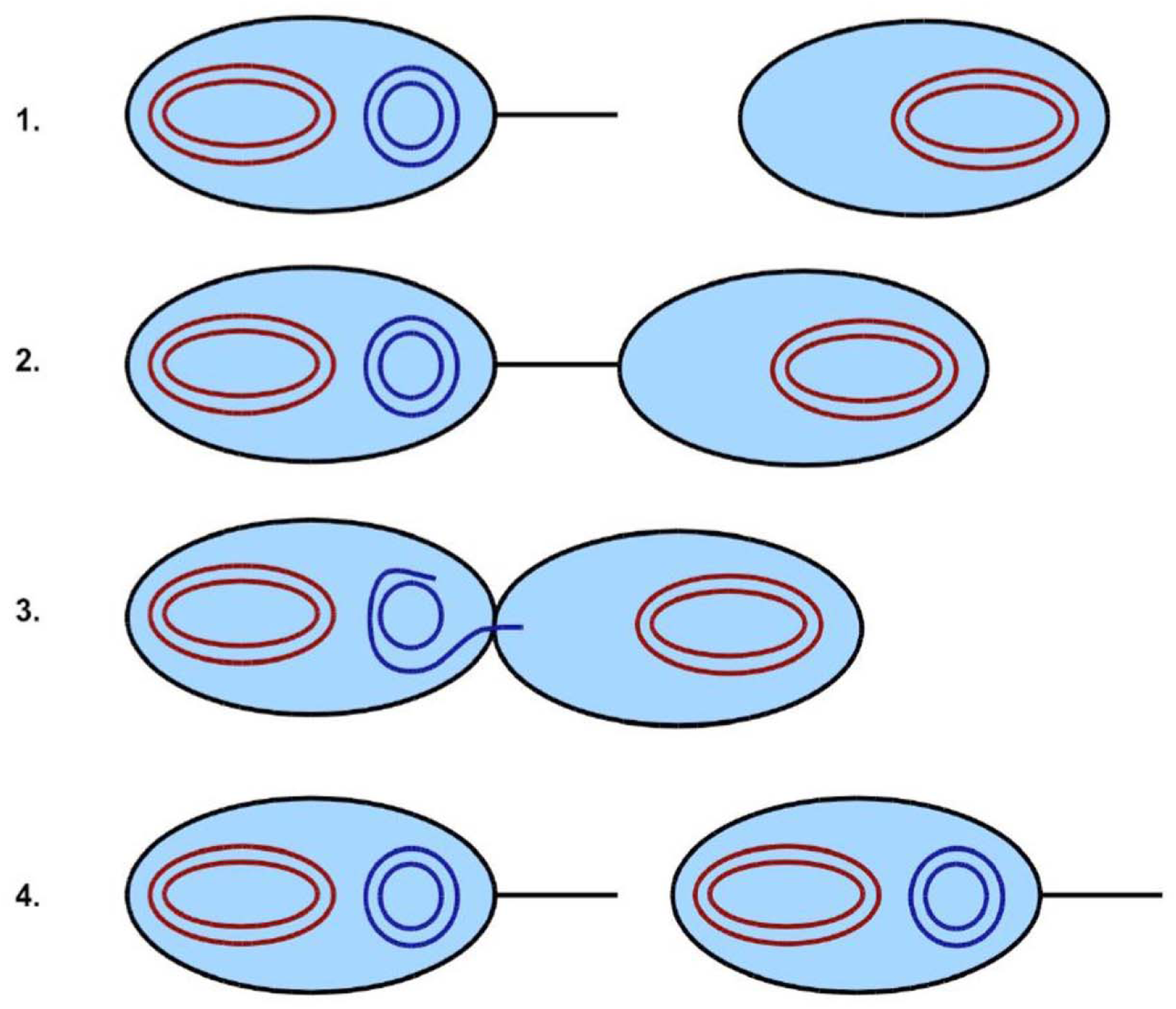
Illustration of the process of bacterial conjugation. In steps 1 and 2, an F ^+^bacterium containing the F-plasmid (blue) binds to an F^−^ bacterium lacking the plasmid. One of the template strands from the F plasmid then moves into the F^−^ bacterium, as shown in step 3. In step 4, the complementary strands are synthesized to reform the complete F-plasmids in both bacteria. Both bacteria are now of the F ^+^ type. Figure 1 was reused with permission from the Genetics Society of America (Raz and Tannenbaum 2010)

However, the F^+^/F^−^ system is in some ways atypical for bacterial conjugation systems, due to its conjugative machinery rapidly switching to a de-repressed state after a conjugation event (Ghigo 2001), while in nature most conjugative plasmid systems are repressed via the expression of T4SS modules (Frost and Koraimann 2010; Andam and Gogarten 2013; Koraimann and Wagner 2014). It is believed the reason behind the propensity of the K-12 strain to switch to the de-repressed state lies in a reduction of the metabolic costs associated with continuously maintaining the enzymatic machinery necessary for conjugation (Ghigo 2001). Indeed, it is known that mutant forms of the Fplasmid system are permanently de-repressed (Ghigo 2001), so in general it is quite possible that these are the strains that were accidentally generated under the experimental conditions that the plasmids were being studied in the experiment which they were originally discovered by Lederberg (Tatum and Lederberg 1947). Consequently, conjugation for the K12 strain occurs much faster than it does in other strains of *Escherichia Coli*.

Nevertheless, the F-plasmid system is still one of the best characterized bacterial conjugation systems, and it can be considered to be broadly representative of most bacterial conjugation systems. Therefore, it makes sense to base mathematical models of conjugation-mediated HGT on the F-plasmid system.

In recent years there has been a steady increase in new theoretical works regarding HGT that included various quasi-species models (Park and Deem 2007; Munoz *et al.* 2008), continuous and stochastic kinetic models that have integrated HGT and investigated its effects on population fitness and its implications on the outspread of novel traits in a population (Nielsen and Townsend 2004; Koslowski and Zehender 2005; Novozhilov *et al.* 2005; Levin and Cornejo 2009; Philipsen *et al.* 2010; Mc Ginty *et al.* 2013), and on the effect of HGT on the distribution of fitness of a bacterial population(Mozhayskiy and Tagkopoulos 2012). However, little effort has been made regarding the evolutionary dynamics of antibiotic resistance through HGT (Maclean *et al.* 2010), with the exclusion of our previous work regarding this subject for a static landscape (Raz and Tannenbaum 2010). We note that while our current model provides the framework for a more in-depth analysis on the implications of horizontal gene transfer on the error-catastrophe phenomenon in a dynamic landscape, we have chosen to deter this analysis for future work.

Recently, the authors have reported an updated mathematical model which has introduced a repression/de-repression mechanism to the framework of our model (Atsmon-Raz and Tannenbaum 2014). This seemingly subtle addition was sufficient to have significant effects on the mean fitness of the simulated bacterial population which were dependent on the rate of repression. More importantly, it introduced a transition value for the repression rate which allows an amelioration of the expected fitness cost due to the maintenance of the antibiotic resistance which was predicted by our model in agreement with other studies (Dahlberg and Chao 2003; Dionisio 2005; Sousa *et al.* 2012; Baltrus 2013). In both our previous studies, our models predicted a slightly deleterious effect on the mean fitness on the population. However, both of these models were based on a single fitness peak approximation of a static fitness landscape and were solved under the assumption of a steady state.

Herein, we explore the hypothesis that conjugation-mediated HGT facilitates adaptation in dynamic environments and extend our previous model by including a periodically shifting single peak landscape and analytically solving our system for different limits while examining the differences between their effective growth rates. This will allow us to evaluate the adaptation time of our population to an antibiotics and the outspread of the population’s resistance to that antibiotic. Although our results are based on the specific details of our mathematical model, they are nevertheless in qualitative agreement with circumstantial evidence regarding the outspread of antibiotic resistance via HGT and expand current understanding of resistance dynamics through HGT (A generalized description of the processes which our model accounts for is detailed in Equation 1).

## MATERIALS AND METHODS

### A. Definitions of the model

We assume that the genome of each bacterium consists of two DNA molecules. The first DNA molecule contains all of the genes necessary for the proper growth and reproduction of the bacterium itself. It corresponds to the large, circular chromosome defining the bacterial genome. We assume moreover that there exists a wild-type genome characterized by a “master” DNA sequence. It is assumed that a bacterium with the master genome has a wild-type fitness, or first-order growth rate constant, given by 1. Such a bacterium is termed viable. Furthermore, we assume that any mutation to the bacterial genome renders the genome defective, so that the bacterium then has a fitness of 0. Bacteria with defective genomes are termed non-viable. This is known as the *singlefitness-peak approximation* in quasispecies theory, and a detailed formulation was previously given in the literature (Tannenbaum and Shakhnovich 2005).

The second DNA molecule is the F plasmid, which holds the necessary information to transcribe the enzymes that provides antibiotic drug resistance to the bacterium. As with the single-fitness-peak approximation made for the bacterial genome, we assume that there is a master sequence for the antibiotic drug resistance gene. If that gene corresponds to a master sequence, then we assume that the bacterium is resistant to the antibiotic. Otherwise, the bacterium is not resistant to the antibiotic, and is assumed to die with a first-order rate constant *Κ_D_*. Furthermore, we assume that only viable bacteria interact with the antibiotic, since non-viable bacteria do not grow and are treated as dead.

Therefore, if the death rate constant *Κ_D_* is sufficiently large, then it is reasonable to assume that all non-resistant genomes are present in negligible amounts in the population, and adaptation only occurs after a peak shift has occurred. This of course necessitates a bacterial population size adapted to the previous peak that is initially sufficiently large to allow it time to adapt to the new peak before dying off.

We assume that replication errors are due to mismatches during daughter strand synthesis that are sub-sequentially fixed in the genome. We let ε denote the mismatch probability per base-pair, so that ε/2 is the probability of making a mismatch, which is fixed as a mutation in the genome. If *L_v_* and *L_r_* denote the lengths of the viability, conjugation, and resistance portions of the genome, respectively, then we have that *p_v_* = (1 – *ε* / 2)*^Lv^* and *p_r_* = (1 – *ε* / 2)*^Lr^*.

Now, by approximating that *L_v_* and *L_r_* become infinite, we may allow the genome length *L* = *L_v_* + *L_r_* to become very large while holding *μ* = *εL* constant. Therefore, defining *α_v_* = *L_v_* / *L*,*α_r_* = *L_r_* / *L*, we then obtain *p_v,r_* = *e*^−*αv,rμ*/2^ (Tannenbaum and Shakhnovich 2005; Madsen *et al.* 2012).

The semiconservative replication of the bacterial genome is not necessarily error-free, so that a given template strand has a probability of producing a daughter genome that is identical to the original parent. We define *p_v_* as the probability of a bacterium’s chromosome template strand to produce a daughter chromosome without introducing any new point-mutations, and *p_r_* as the probability of the resistance gene in the plasmid to replicate without introducing any new mutations. If we assume that sequence lengths are long, then making an assumption known as the *neglect of backmutations* (Tannenbaum and Shakhnovich 2005), we assume that a template strand derived from a parent that differs from the master genome produces a daughter that differs from the master genome with probability 1. We also assume that there is a probability *p*_01_ that a template strand from the current antibiotic resistance master sequence will produce a daughter that corresponds to the next antibiotic resistance master sequence (the next antibiotic resistance master sequence template strand has an equal probability of producing a current antibiotic resistance master sequence).

We note that unlike our previous model (Atsmon-Raz and Tannenbaum 2014), here we assume that at any given time, all of the bacteria host a plasmid, and that their ability to conjugate (or not) is determined strictly by if they are in a repressed or de-repressed state. In order to model the repression/de-repression dynamics, we will assume that the transition from the repressed to the de-repressed state is characterized by a first-order rate constant k_−+_, and the transition from the de-repressed to the repressed state is characterized by a first-order rate constant k_+−_.

We model the conjugation process by assuming that conjugation may only occur between a viable F^+^ bacterium and a viable F^−^ bacterium. Thus, conjugation can only occur between a bacterium from the *n*_+ *j*_ (*j* = 0 / 1 / −) types and bacterium from the *n*_− *j*_ (*j* = 0 / 1 / −) types. This process is modeled as a second-order collision reaction with a rate constant γ, which is also used in our equations in order to allow for errors in the conjugation and resistance genes that may be caused by faulty horizontal or vertical gene transfer. The conjugation process itself involves the transfer of one of the strands of the plasmid from the F^+^ bacterium to the F^−^bacterium, so that the full plasmid needs to be resynthesized in both bacteria via daughter strand synthesis. We also assume that the system volume changes so as to maintain a constant population density ρ.

It should be emphasized that we are assuming for simplicity that all bacteria in the population contain exactly one plasmid. We also assume that, during conjugation, the plasmid transferred from the F^+^-bacterium replaces the plasmid in the F^−^-bacterium. This is a simplifying assumption that will obviously have to be re-examined in future research, where we anticipate developing more accurate models that allow for variable plasmid numbers in the bacterial cell. The basis for this assumption derives from the observation that plasmids of similar compatibility classes cannot co-exist in the same cell (Novick 1987; Norman *et al.* 2009), and that bacteria can control the number of plasmids in the cell (Uhlin and Nordstrom 1975; Novick 1987; Park *et al.* 2001).

We introduce a dynamic landscape into our model which replaces the previously applied static landscape (Raz and Tannenbaum 2010; Atsmon-Raz and Tannenbaum 2014). We proceed by assuming that after every time τ, the antibiotic “master” sequence shifts by a single point-mutation. We note that in previous work, a similar landscape was investigated in order to study the effect of mutators on time varying fitness landscapes (Gorodetsky and Tannenbaum 2008). To model the dynamics, we assume that *Κ_D_* is sufficiently large, and the point-mutation rate is sufficiently low, that we need only consider bacterial genomes whose antibiotic resistance region differs by at most one point mutation from the given “master” sequence. This assumption is analogous to the one made by Nilsson and Snoad when studying the error threshold in dynamic fitness landscapes (Nilsson and Snoad 2000). However, our application of a dynamic landscape differs from the Nilsson and Snoad implementation in two key points: 1) After a peak shift, they used a populations permutation vector instead of monitoring the actual adaptive dynamics to the next peak or in other words, after a peak shift, there was already a small fraction of organisms that were adapted to the new peak, which then drove the growth of the population. 2) In the Nilsson and Snoad model, both viable and unviable organisms were able to grow with the viable organisms being assigned a significantly faster growth rate over the unviable organisms, while in our model we will consider only viable bacteria.

### B. Evolutionary Dynamics Equations

To derive the evolutionary dynamics of our dynamic landscape, we make a few additional assumptions beyond what was made in our previous work. First of all, we assume that the length of the genome controlling conjugation is sufficiently short that we may neglect any mutations to it, so that all F- plasmid in the population are functional. What determines whether a bacterium is an F+ bacterium or an F- bacterium depends only on whether the plasmid is repressed or de-repressed. We then consider the dynamics from immediately before a peak shift to immediately before the following peak shift. We denote our various sub-populations by *n*_±0/1/−_ where the first index is a “+” if the organism has a de-repressed plasmid (i.e., it is F^+^), while the index is a “-“ if the organism has a repressed plasmid (i.e. it is F^−^). The second index is a “0” if the organism is resistant to the antibiotic immediately before the peak shift, while the index is a “1” if the organism becomes resistant to the antibiotic after the peak shift. The same index can also be “-“ if the organism is not resistant to the current or the next antibiotic.

Therefore, allowing us to summarize all of the processes that are relevant for each subpopulation in our model into a generalized list of balance equations which are schematically illustrated in Figure 2 and listed in as reaction equations in Equation (1).

**Fig 2.**
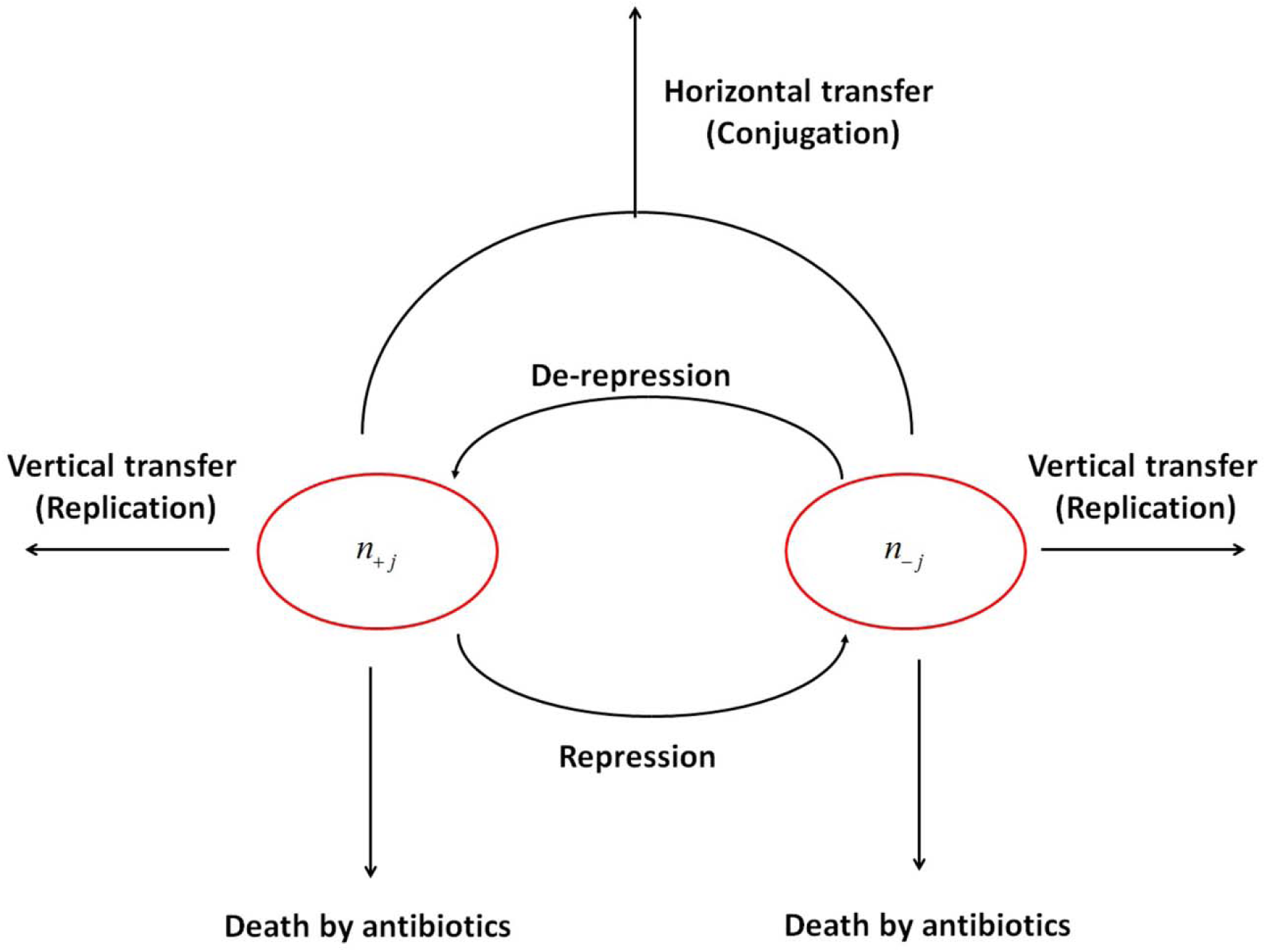
Schematic of the main evolutionary processes which are considered within the framework of our model. We note that both vertical and horizontal transfer confer a probability of mutation.

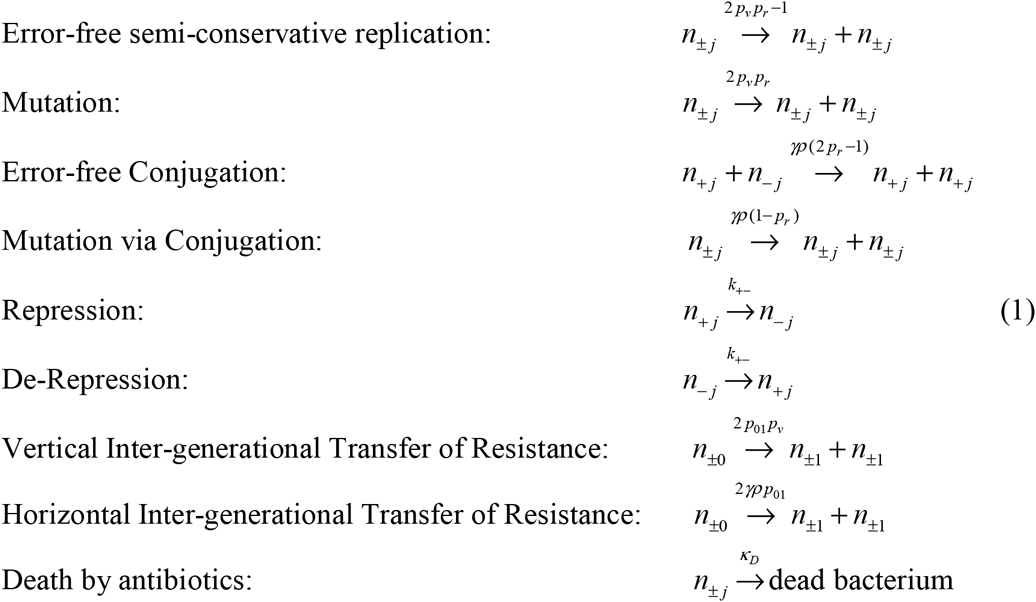

We note that *p_j_* denotes two possible values for the resistance gene – 0 and 1. 0 denotes resistance to the active antibiotic in the current cycle and 1 denotes resistance for a different antibiotic that will be presented to the population in the next cycle.

If we use the notations 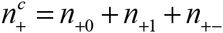 and 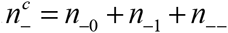, then the evolutionary dynamics equations immediately following the peak shift can be described as follows,

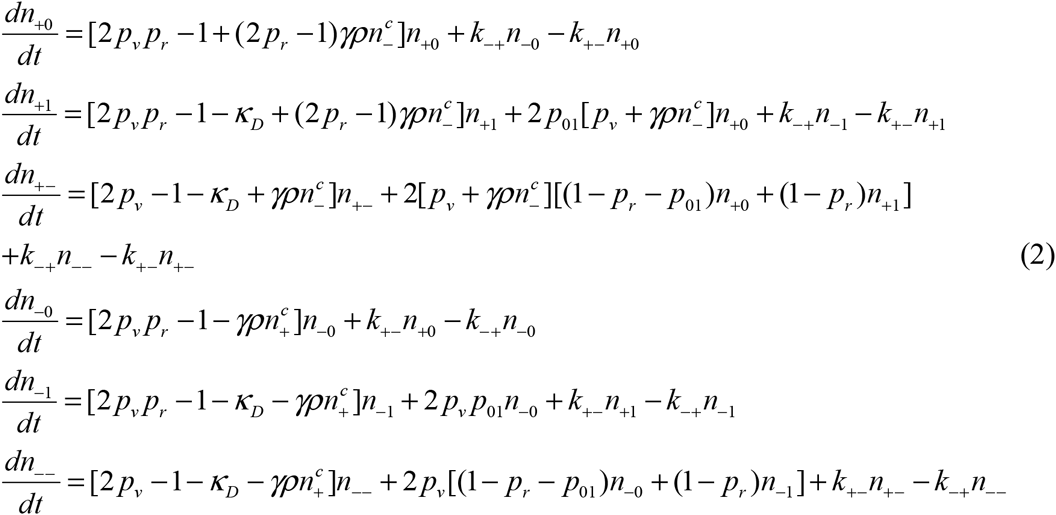

In the following sections we detail the analytical solutions for this system of equations under various conditions. We have tested numerically the results and approximations in these sections with the Matlab 2012b ode45 function and visualized it with the in-built plot function. Furthermore, we have validated our analytical solutions for the integration, derivation and approximations of these equations with Wolfram Mathematica 10.

## RESULTS

### A. Derivation of the condition for adaptability for *γρ* → 0, finite *Κ_D_* limit

When *γρ* = 0, we have the following:

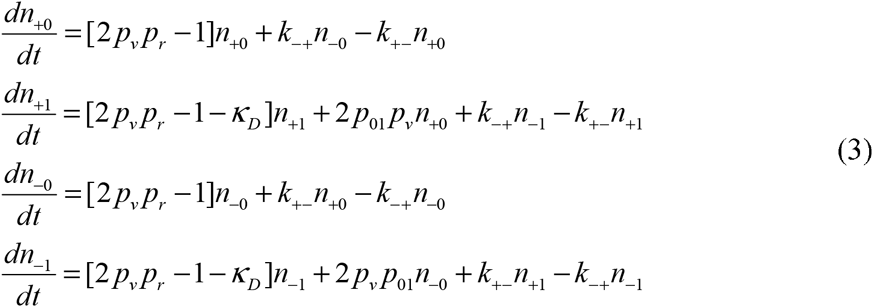

Defining *n*_⏢_ = *n*_+⏢_ + *n*_−⏢_, we obtain,

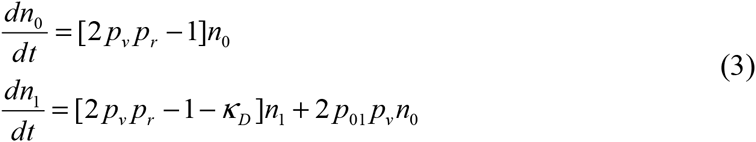

Integration of these equations will give us the following terms which we have plotted in Figure 3:

**Figure 3.**
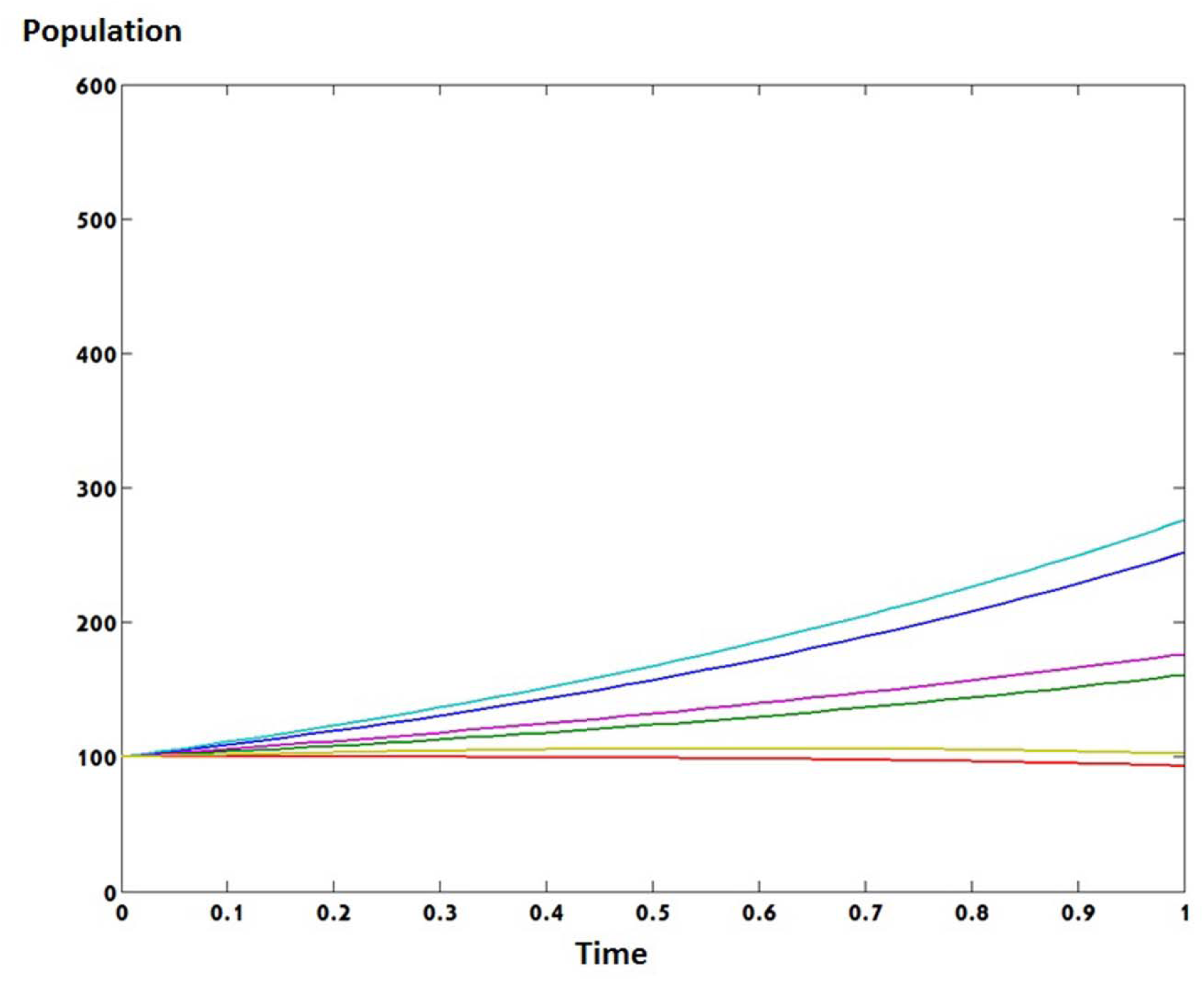
Numerical solution for *γρ* → 0, finite *Κ_D_* limit. The blue line is for *n*+0, the green line is for *n*_+1_, the red line is *n*_+−_, the cyan line is *n*_?0_, the purple line is *n*_−1_ and the yellow line is *n*_−−_. Chosen parameters were *p_v_* = 0.99; *p_r_* = 0.995;*Κ_D_* = 0.7;*γρ* = 0; *k*_−+_ = 0.8; *k*_+−_ = 0.9; *p*_01_ = 0.1

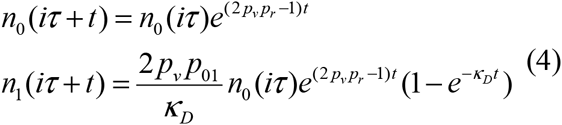

Since *Κ_D_* is assumed to be large, the criterion for adaptability is that the viable, resistant population grows faster than the viable non-resistant population over a cycle. We denote the ratio between these populations as *ϕ* and require this ratio to be greater than 1 so that,

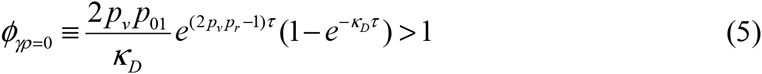

If we define 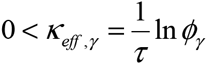, then the criterion for adaptability becomes,

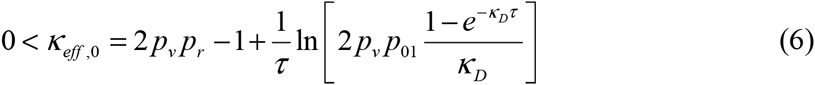

### B. Derivation of the condition for adaptability for *γρ* → ∞, finite *Κ_D_* limit

When *γρ* is large, we may assume that *n*_−0_, *n*_−1_, *n*_−_ → 0. We may also assume that *γρn*_−0_, *γρn*_−1_, *γρn*_−−_ converge to well-defined, finite values. We can then take the terms of 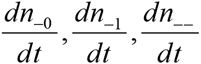 from Equation 2 and by assuming that they reach a steady state we have,

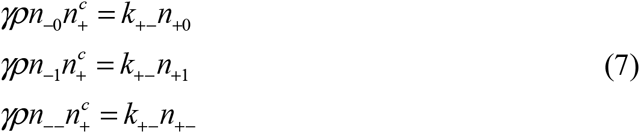

Summing these equations gives,

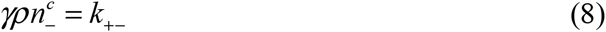

And so the dynamical equations become,

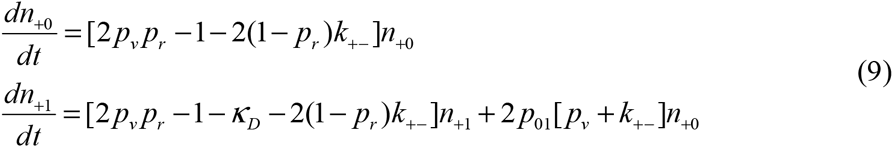

We can use the analytical solution (Section A of the SI) for these differentials equations to formulate the criterion for the adaptability for large *γρ* following a similar procedure to that of the previous subsection, which gives us the term,

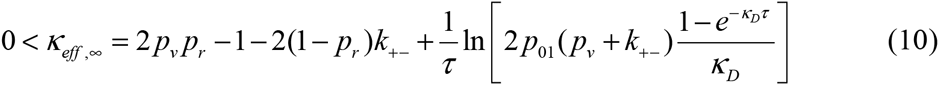

and so note that,

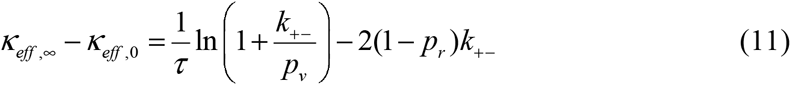

Furthermore, we can equate the terms for *n*_+0_ and *n*_+1_ and find that *n*_+0_ competes with *n*_+1_ and eventually exceeds its population at a given time (Figure 4):

**Fig 4.**
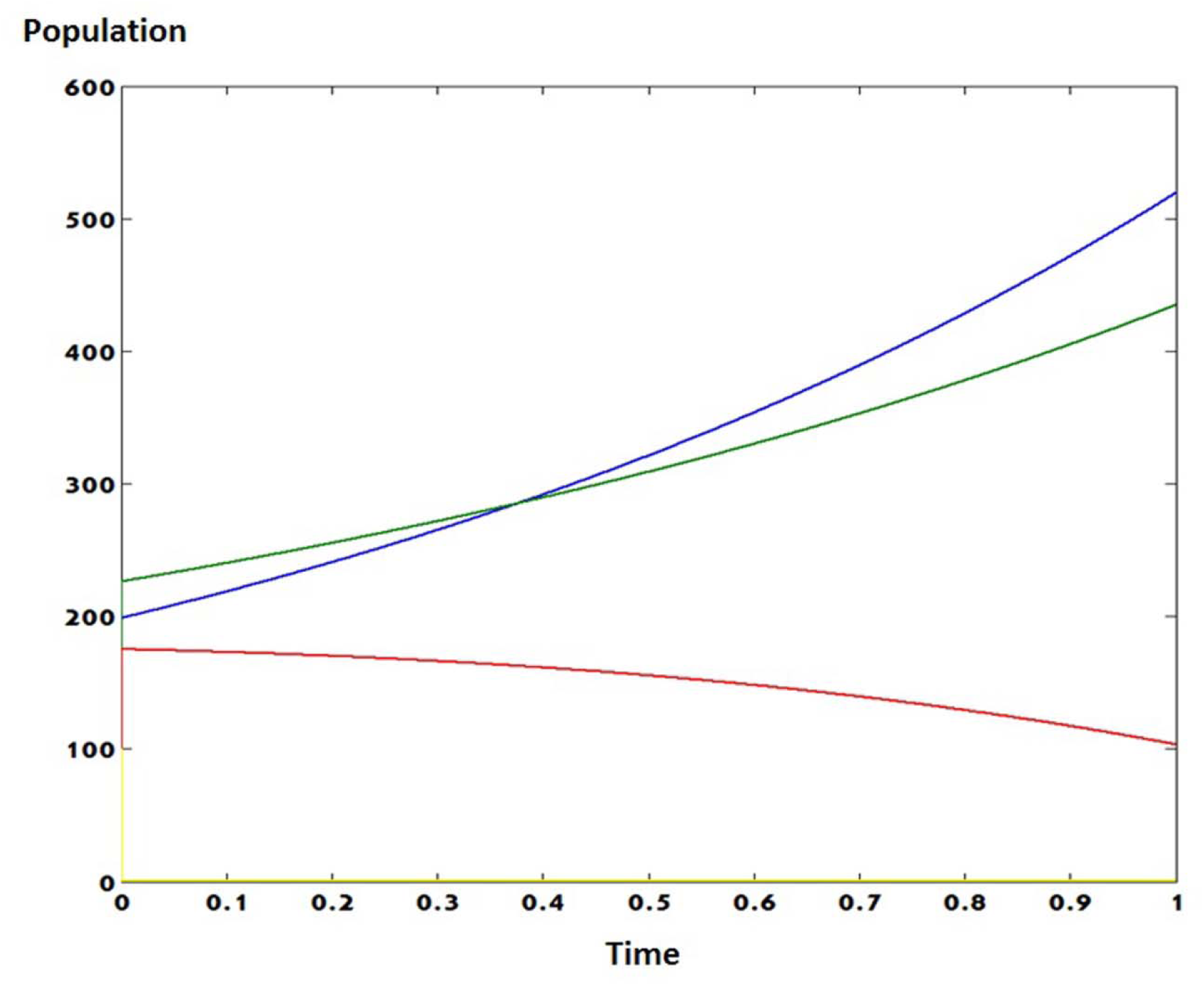
Numerical solution for *γρ* → ∞, finite *Κ_D_* limit. The blue line is for *n*_+0_, the green line is for *n*_+1_, the red line is *n*_+−_, the cyan line is *n*_−0_, the purple line is *n*−1 and the yellow line is *n*_−−_. Chosen parameters were *p_v_* = 0.99; *p_r_* = 0.995;*Κ_D_* = 0.7;*γρ* = 1000; *k*_−+_ = 0.8; *k*_+−_ = 0.9; *p*_01_ = 0.1

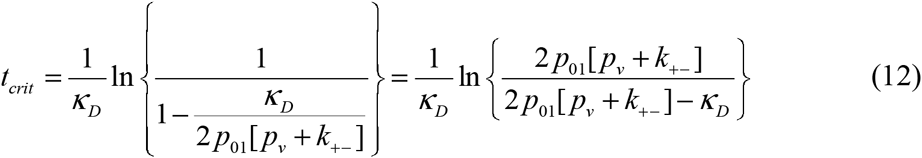

### C. Derivation of the condition for adaptability for the *Κ_D_* →∞, finite*γρ* limit

When *ΚD* is large, we may assume that 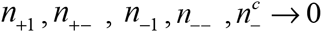 We may also assume that *Κ_D_*^*n*^_+1_, *Κ_D_*^*n*^_+−_, *Κ_D_*^*n*^_−1_ and *Κ_D_*^*n*^_−−_ converge to well-defined, finite values.

We can then sum the terms for *n*_±0_ and *n*_±1_ in order to derive:

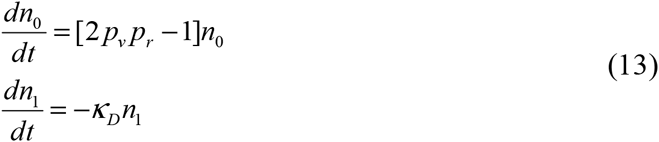

We note that a plot of all of the system’s sub-populations has been placed in Figure 5.

**Fig 5.**
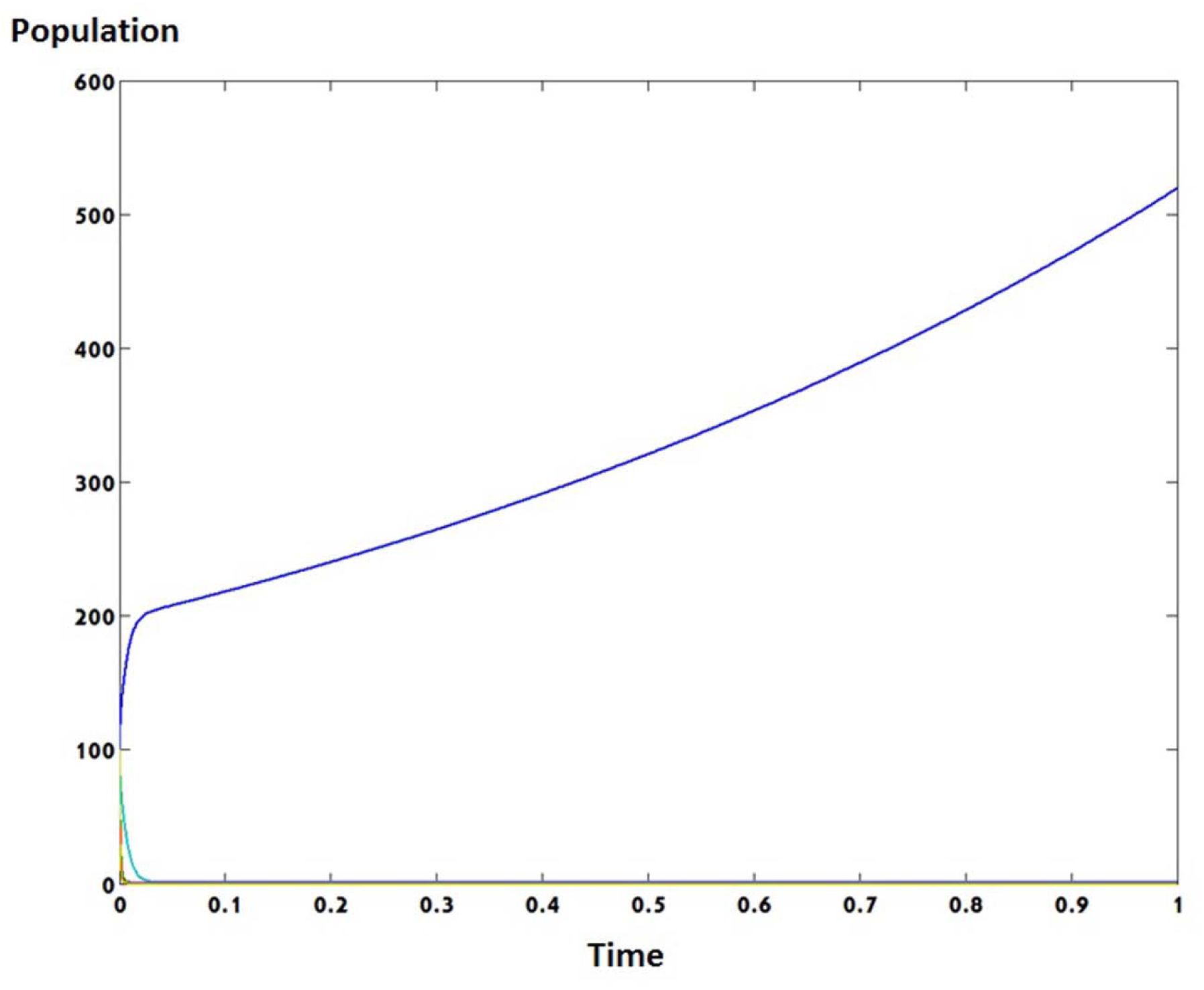
Numerical solution for *Κ_D_* →∞, finite *γρ* limit. The blue line is for *n*_+0_, the green line is for *n*_+1_, the red line is *n*_+–_, the cyan line is *n*_−0_, the purple line is *n*−1 and the yellow line is *n*_−−_. Chosen parameters were *p_v_* = 0.99; *p_r_* = 0.995;*Κ_D_* = 1000;*γρ* = 0.9; *k*_−+_ = 0.8; *k*_+–_ = 0.9; *p*_01_ = 0.1

By following a similar process to the one which we described in the previous sections we may calculate the term for *ϕ* which takes the form of

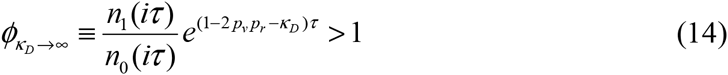

However, the terms for *n*_0_ and *n*_1_ in this case are independent of each other’s initial populations and the resulting term for *Κ*_*eff*, *Κ* →∞_ is

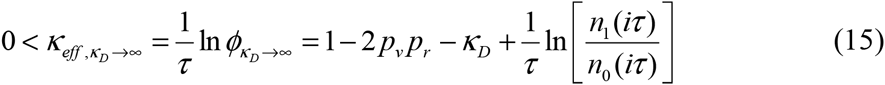

### D. Derivation of the condition for adaptability for the *Κ_D_* → ∞,*γρ* →∞ limit

In order to find a solution for our system within these limits, we may investigate the limiting case of *Κ_D_* → ∞ for the solution of *n*_+0_ and *n*_+1_ (as detailed in Section A of the SI).

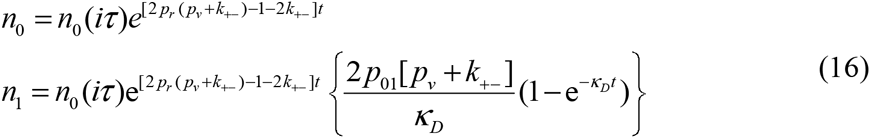

As in the previous sections, a plot of all of the system’s sub-populations is illustrated in Figure 6. Furthermore, since *Κ_D_* → ∞, *n*_1_ takes the following form

**Fig 6.**
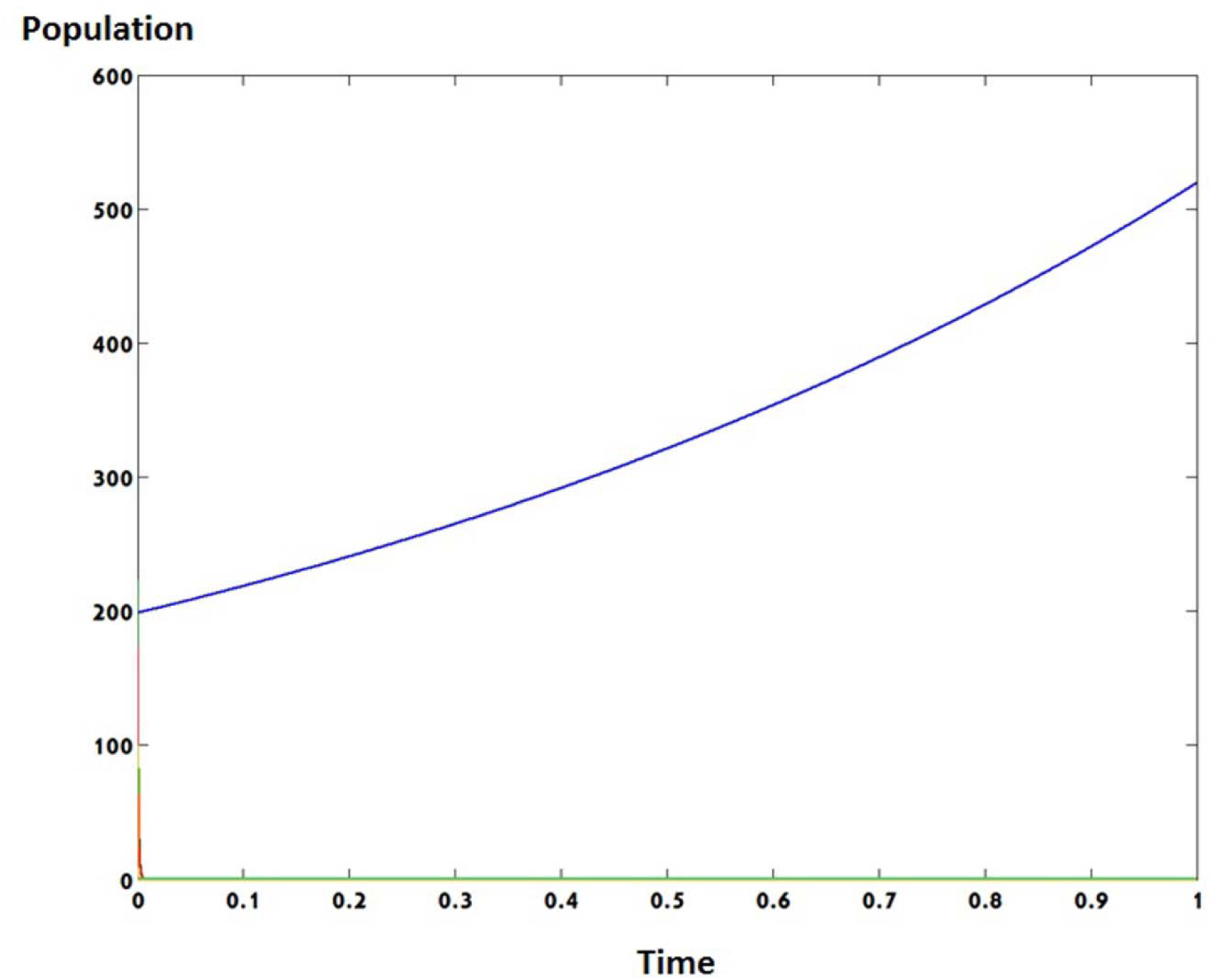
Numerical solution for *Κ_D_* → ∞,*γρ* →∞. The blue line is for *n*_+0_, the green line is for *n*_+1_, the red line is *n*_+–_, the cyan line is *n*_−0_, the purple line is *n*_−1_ and the yellow line is *n*_−−_. Chosen parameters were *p_v_* = 0.99; *p_r_* = 0.995;*Κ_D_* = 1000;*γρ* = 1000; *k*_−+_ = 0.8; *k*_+–_ = 0.9; *p*_01_ = 0.1

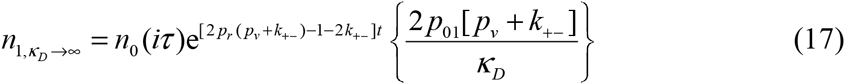

While *n*_1_ maintains its original form as in Equation 9.

Therefore, the effective growth rate can be by applying the *Κ_D_* →∞ limit to the *Κ*_*eff*, *γρ*→∞:_

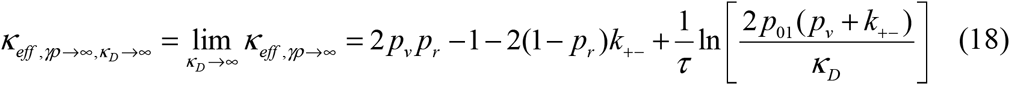

We have summarized the effective growth rates for each limit case in Table 1.

**Table 1.** Effective growth rates for different limit cases of our model.

| Limits | $\kappa_{eff}$ |
| --- | --- |
| $\gamma p \rightarrow 0$ | $2p_v p_r - 1 + \frac{1}{\tau} \ln \left[ 2p_v p_{01} \frac{1 - e^{-\kappa_D \tau}}{\kappa_D} \right]$ |
| $\gamma p \rightarrow \infty$ | $2p_v p_r - 1 - 2(1 - p_r)k_{+-} + \frac{1}{\tau} \ln \left[ 2p_{01}(p_v + k_{+-}) \frac{1 - e^{-\kappa_D \tau}}{\kappa_D} \right]$ |
| $\kappa_D \rightarrow \infty$ | $1 - 2p_v p_r - \kappa_D + \frac{1}{\tau} \ln \left[ \frac{n_1(i\tau)}{n_0(i\tau)} \right]$ |
| $\kappa_D \rightarrow \infty, \gamma p \rightarrow \infty$ | $2p_v p_r - 1 - 2(1 - p_r)k_{+-} + \frac{1}{\tau} \ln \left[ \frac{2p_{01}(p_v + k_{+-})}{\kappa_D} \right]$ |

### E. Non conjugator to conjugator transition for condition for adaptability for *k*−+ = 0

When *k*_−+_ > 0, then the population will always consist of a certain fraction of conjugators, no matter what the value of *γρ*. However, as *k*_−+_ approaches 0, the fraction of conjugators may exhibit more complex behavior. Here, the production of conjugators is driven entirely by the conjugation process itself, whose rate is characterized by the parameter *γρ*. The inhibition of conjugators, on the other hand, is characterized by a rate constant *k*_+–_. Presumably, the relate values of *γρ* versus *k*_+−_ will determine the fraction of conjugators in the population.

In particular, if we define *λ* = *γρ*, then we wish to determine the critical value of *λ*, denoted as *λ_c_*, which below its value there are no conjugators in the population. So, let us define *n*_⏧⏧,0_ = (*n*_⏧ ⏧_)_*λ* =*λ*_, and let us define *n*_⏧ ⏧,1_ = (∂*n*_⏧ ⏧_ / ∂*λ*)_*λ* =*λ c*_ Using the fact that there are no conjugators (*n*_+0, 0,_ *n*_+1,0,_ *n*_+<−,0,_ 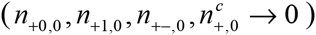 when *λ* = *λ_c_*, we have that,

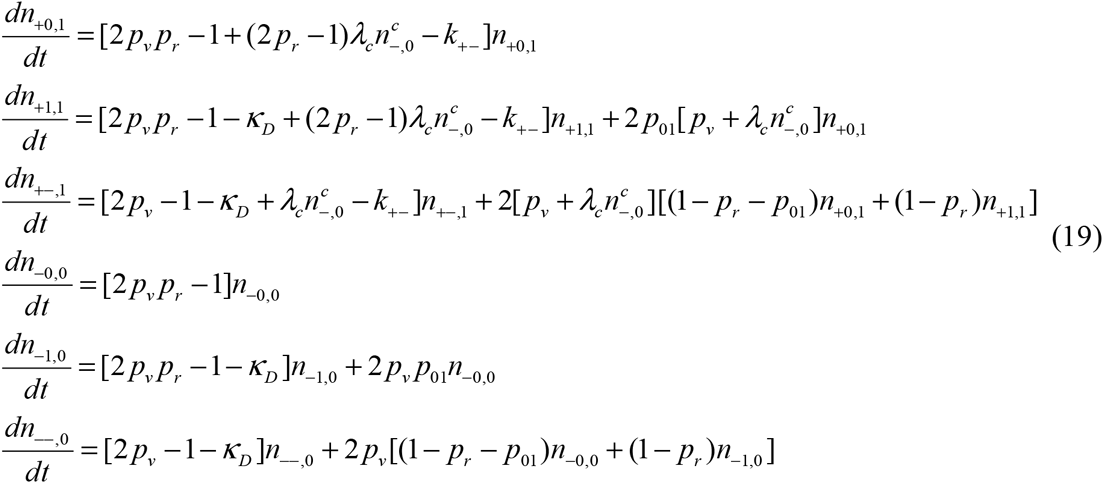

To lowest order in *λ* –*λ_c_* (i.e. zeroth order), we then have that the effective growth rate constant of the non-conjugator population is given by

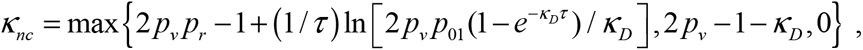

where the first term holds if the resistant non-conjugator population adapts; the second term holds if the resistant, non-conjugator population does not adapt, but the viable, nonresistant population still has a positive growth rate; and the third term holds if the viable, non-conjugator population has a negative first-order growth rate constant.

For the conjugators to begin to rise to a positive fraction of the population once *λ*exceeds λ_*c*_,we must have that the effective growth rate of the conjugators exceeds the effective growth rate of the non-conjugator population.

While the effective growth rate constant of the resistant conjugators at *λ* = *λ_c_* is given by,

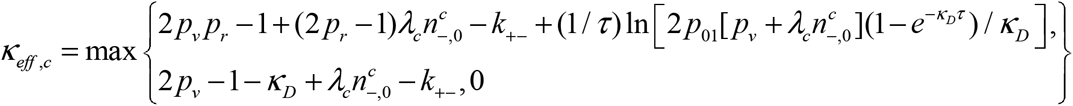

The first solution holds if the resistant conjugator population adapts to the new antibiotic, the second expression holds if the resistant conjugator population does not adapt but the viable population has a positive growth rate. We get the third possibility 0 if the viable conjugator population has a negative first order growth rate constant.

Therefore, finding the transition value of *λ_c_* is dependent on the adaptation of both the conjugator and non-conjugator populations to the antibiotic.

We can find the transition value of *λ_c_* for the non-conjugator to conjugator value by equating *Κeff*, *c* with *Κ*_*eff*, *nc*_. However, we need to consider two cases: 1) When all conjugators (repressed and de-repressed) have adapted to the antibiotic (detailed in section B of the SI); 2) When all conjugators (repressed and de-repressed) have not adapted to the antibiotic yet they maintain a positive effective growth rate (detailed in section C of the SI).

For the first case

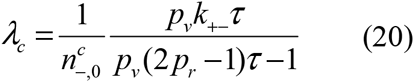

For the second case

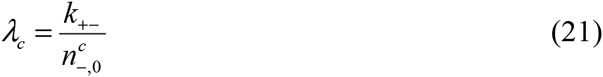

We have analytically solved the term for 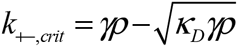 and detailed it in section D of the SI.

### F. Repression driven restoration of fitness for the limit cases of *γρ* → ∞ and for *Κ_D_* → ∞,*γρ* →∞

In our recent study, we have investigated a similar model for a static landscape and found that the fitness cost that antibiotic resistance has on population of prokaryotic bacteria maybe restored via repression when the value of the repression reaches a critical value.(Atsmon-Raz and Tannenbaum 2014) Herein, we were interested if our current system would also demonstrate such a behavior and indeed identified such a critical local maximum of *k*_+–_ for both the limiting cases of *γρ* → ∞ and for *Κ_D_* → ∞,*γρ* →∞ (SI, Section E):

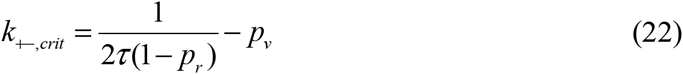

When we examine the possible values for *k*_+–_ we found that when *k*_+–_ > *k*_+–_,_*crit*_, repression is fast enough to decrease the total number of available conjugators, therefore leading to a decrease in the effective growth rate. However, when 0 < *k*_+–_ < *k*_+–_,*crit*, repression assists in the restoration of fitness that would otherwise be lost from the maintenance of antibiotic resistance (Figure 7).We have evaluated the expected effect that repression has on the effective growth rate in section F of the SI.

**Fig 7.**
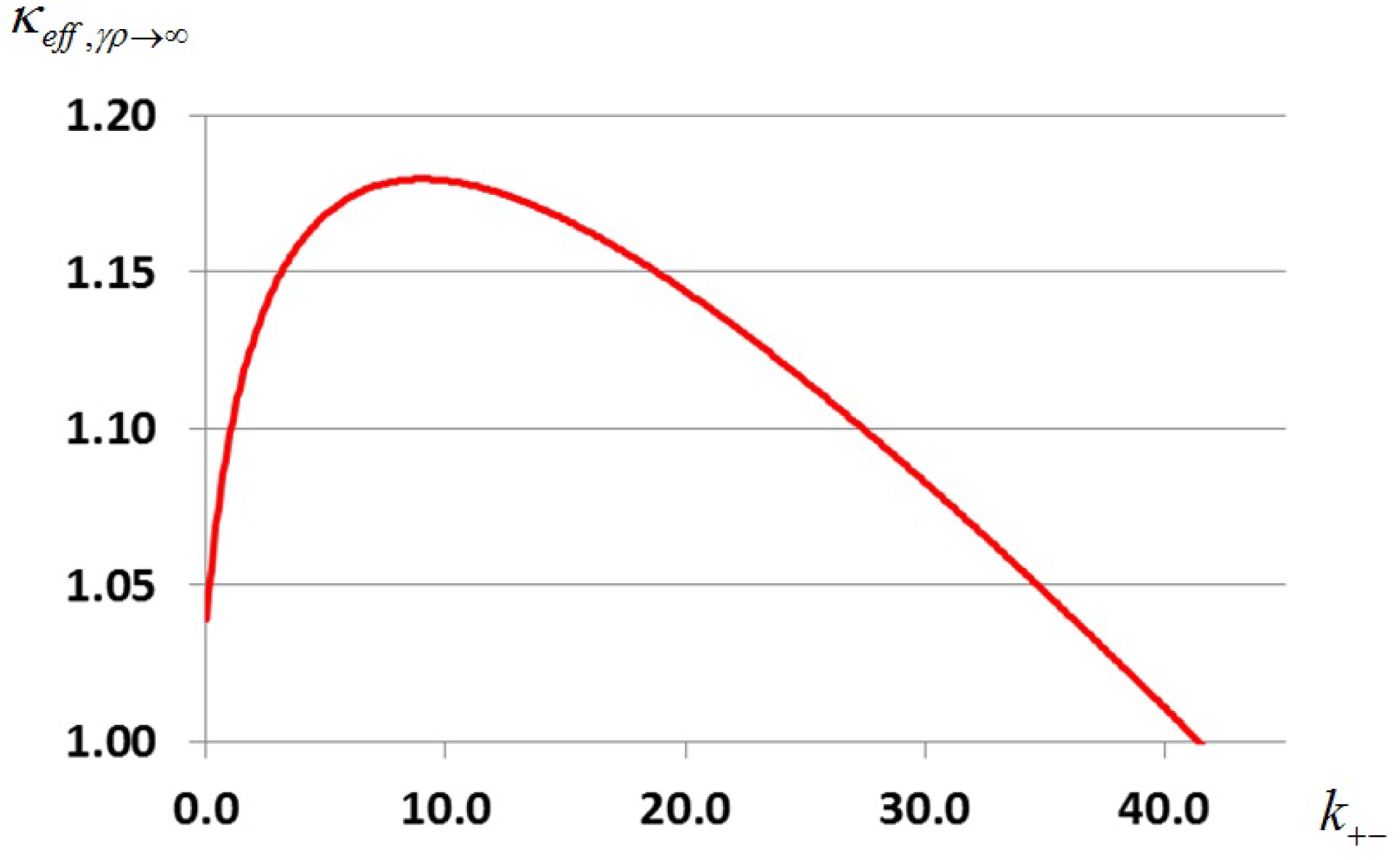
Plot of the effective growth rate *Κ_eff_* vs *k*_+–_ for the limit case of *γρ* → ∞, finite *Κ_D_* limit. Chosen parameters were *p_v_* = 0.99; *p_r_* = 0.995;*Κ_D_* = 0.7; *p*_01_ = 0.1

### G. Negative to Positive transition times for the limits of *Κ*_*eff*, *γρ* →0_, *Κ*_*eff*, *γ* →∞_, and *Κ eff*, *Κ D* →∞, *γρ* →∞

In order to find how long it would take a bacterial population to adapt to a new antibiotic we examined the conditions of adaptability for each limit case and equated them to zero to find the value of the expected adaptation times for each case. Detailed solutions can be found in Sections G-I of the SI) and compared their values by setting the values of all parameters so that *p_v_*, *p_r_*, *p*_01_,*Κ_D_*, *k*_+–_ →1 We have summarized these results into Table 2 and will discuss them in detail in the following section.

**Table 2.** Negative to positive transition times for our studied limit cases.

| Limits | $t_{crit}$ | Approximated value |
| --- | --- | --- |
| $\gamma p \rightarrow 0^*$ | $\frac{-\alpha + \sqrt{\alpha^2 + 4\alpha\{2p_v p_r - 1 - 2(1 - p_r)k_{+-}\}}}{2\alpha\{2p_v p_r - 1 - 2(1 - p_r)k_{+-}\}}$ $* \alpha = 2p_{01}(p_v + k_{+-})$ | $\frac{-1 \pm \sqrt{1+2}}{2} = 0.366$ |
| $\gamma p \rightarrow \infty$ | $\frac{-1 + \sqrt{1 + 4(2p_v p_r - 1)\left(\frac{\kappa_D}{2p_v p_{01}}\right)}}{2(2p_v p_r - 1)}$ | $\frac{-1 + \sqrt{2}}{2} = 0.207$ |
| $\kappa_D \rightarrow \infty, \gamma p \rightarrow \infty$ | $\frac{\ln\left[\frac{2p_{01}(p_v + k_{+-})}{\kappa_D}\right]}{2(1 - p_r)k_{+-} - 2p_v p_r - 1}$ | $\frac{\ln(4)}{3} = 0.462$ |

## DISCUSSION AND CONCLUSIONS

In the last two decades, numerous reports have detailed the emergence and rapid outspread of antibiotic resistance in bacterial populations (Neu 1992; Davies and Davies 2010; French 2010a; Roca *et al.* 2015). Several well studied examples of antibiotic drug resistance include penicillin (Tatum and Lederberg 1947), methicillin (Grundmann *et al.* 2006), tetracycline (Speer *et al.* 1992), fluoroquinolones (Tapsall and Organization 2001), cephalosporins (Livermore 1987), vancomycin (Hiramatsu 1998) and carbapenem (Bratu *et al.* 2005; Sahuquillo-Arce *et al.* 2014). Furthermore, these resistances may combine into new strains that are resistant to multiple types of antibiotics for a bacterial population (Dijkshoorn *et al.* 2007; Boucher *et al.* 2009; Fischbach and Walsh 2009; Magiorakos *et al.* 2012). Consequently, the threat imposed by the growing outspread and evolution of antibiotic drug resistance may be considered to be of severe magnitude when we take into account the current lack of development in the field of novel antimicrobial drugs due to the high costs entailed in their development (Norrby *et al.* 2005; Spellberg *et al.* 2008). Therefore, it has become essential to seek out new strategies to inhibit or even divert the evolutionary path that leads to the rapid adaptation of bacterial populations to novel antibiotics.

Bacterial-conjugation mediated HGT is used by bacterial populations to allow a more rapid adaptation process to environmental changes such as novel antibiotics. Since its discovery (Tatum and Lederberg 1947), it was found to play a critical role in the transmission of antibiotic resistance in prokaryotic populations (Gogarten *et al.* 2002; Osborn and Boltner 2002; Grohmann *et al.* 2003; Burrus and Waldor 2004; Koslowski and Zehender 2005; Novozhilov *et al.* 2005; Bennett 2008), in the formation of biofilms (Drenkard and Ausubel 2002; Madsen *et al.* 2012), and in the transmission of genetic information between bacteria and fungi (Heinemann and Sprague 1989). More recent studies have suggested that HGT-related resistance is expected to overlap on a molecular level with chemotherapy resistance in cancerous tumors (Lambert *et al.* 2011; Trejo-Becerril *et al.* 2012).

From an evolutionary perspective, elevated HGT levels can be considered to be an analogous adaptation strategy to the SOS response in prokaryotes (Metzgar and Wills 2000; Matic 2013) due to the complimentary strand synthesis of the plasmid that occurs in bacterial conjugation. However, while the increased mutation rate in the SOS response is caused by Pol II/IV/V (Cirz *et al.* 2005), in HGT it can be caused by an increase in the conjugation rate itself (Beaber *et al.* 2004). This implies on the importance of regulation in HGT just as is the case for hyper-mutation (Denamur and Matic 2006). In both mechanisms, adapting to a new environment (i.e., antibiotics), confers a fitness cost. In bacterial conjugation-mediated HGT, this cost originates from the addition and integration of the plasmid within the hosting bacterium due to the extended demand for the bacterium’s enzyme machinery, nucleotides, amino acids (for the transcription of the conjugation and resistance related proteins) and any other resources which are required by the plasmid. However, the extent of this cost, and the bacterium’s ability to compensate for it is widely debated (Anderson 1968; Baquero and Blázquez 1997; Levin 2001; Elena and Lenski 2003; Andersson and Hughes 2010; Maclean *et al.* 2010; Perron *et al.* 2010; Baltrus 2013). Previous studies have reported that in some bacteria, developing resistances to specific antibiotics such as VanA (i.e., vancomycin) entails a high fitness cost which could be exploited in future design of therapies (Foucault *et al.* 2009; Andersson and Hughes 2010). While these fitness costs can be ameliorated through compensatory mutation (Lawrence and Ochman 1997; Dahlberg and Chao 2003; Perron *et al.* 2010), it has only been recently suggested that an alternative strategy exists by the induction of the repression mechanism (Atsmon-Raz and Tannenbaum 2014).

Herein, we have described and analyzed a mathematical model that made use of the quasispecies framework as was originally presented by Eigen (Eigen 1971) and later extended by Shakhnovich (Tannenbaum and Shakhnovich 2005) to describe a population of unicellular bacteria that are forced to periodically adapt via time varying fitness landscape to antibiotics. We have allowed this population to use bacterial conjugation mediated-horizontal gene transfer which was regulated via repression and de-repression. Our purpose in this model’s construction was to gain new insights into the dynamics of antibiotic resistance and fitness cost amelioration via HGT.

We analytically solved the effective growth rates of the model for four different limit cases *γρ* →0, *γρ* → ∞, *Κ_D_* → ∞ and *Κ_D_* → ∞,*γρ* →∞. The obtained results have provided us unique terms that allowed us to evaluate the time it would take for the effective growth rate to switch from a negative value (due to the exposure to the new antibiotics) to a positive value (when the population has overcome the evolutionary obstacle presented by the antibiotic), which essentially can be described as the adaptation time of the population after a shift of the fitness landscape. We evaluated the terms for the transition time for each of the limit cases and found that HGT allows a quicker adaptation time for the population then vertical evolution which falls in qualitative agreement with previously published results (Cohen *et al.* 2005; Park and Deem 2007). Interestingly, the adaptation time for the *γρ* → ∞,*Κ_D_* →∞ case was actually found to be slower than for the *γρ* → 0 and the *γρ* →∞ cases. This is likely caused by the elevated level of antibiotics which effectively decreases the pool of potentially resistant bacteria which makes it much harder for the resistance to fixate in the population. Consequently, the adaptation time to the new antibiotic that is required by the population is longer than the *γρ* → 0 and the *γρ* →∞ cases. For the reader’s convenience, we have summarized the effective growth rates and the positive-to-negative transition times respectively, in Table 1 and Table S1. Furthermore, from a pharma-kinetic perspective, these results can be used to evaluate the relevant timescales that would be required for a sequential multidrug strategy to succeed if it was aimed at out-competing the adaptation rates of a bacterial population (Maclean *et al.* 2010).

As in our previous work regarding a static landscape (Atsmon-Raz and Tannenbaum 2014), we have observed a phase transition in the value of the repression constant (*k*_+–_) that held for both the cases of *γρ* →∞ and *γρ* → ∞,*Κ_D_* →∞. However, in our previous work the critical value of *k*_+–_ was determined to be a local minima of the population mean fitness and dependent only on the values of the conjugation (*γρ*) and the antibiotics death rate (*Κ_D_*) with a dependence of 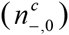. Here, we have found *k*+–,*crit* to be inversely dependent on the probability of correct plasmid replication and the cycle time between peak shifts (*τ*) as well as linearly dependent on the probability of correct replication of the bacterium’s genome (Equation 22).

We can rationalize the dependence on *τ* from the dynamic nature of this model which necessitates that the repression rate has to be limited by the time scale of a peak shift in order for it to have any significance on the existing populations within any given cycle. In the case of the static landscape model, due to its independence of time, such a limitation is irrelevant. We may also rationalize the absence of the conjugation rate value (*γρ*) in our current model, since our term for *k*+–,*crit* in our current model, is derived directly from the approximated terms of the effective growth rates for the limit cases of *γρ* →∞ and *γρ* → ∞,*Κ_D_* →∞ while in our previous model it was derived directly from the mean fitness function which mathematically differs from the formulation of the effective growth rates. However, at this point we are unable to rationalize the absence of *Κ_D_* nor the presence of the errorless replication probabilities in our new term for *k*_+–,*crit*_.

Further analysis of the *k*+–,*crit* term has revealed that it is a local maximum of the effective growth rate *Κ_eff_* (Section E of the SI). Therefore, we can split the effect that *k*_+–_ has on *Κ_eff_* into two regions which define two different “behaviors” for our system. The first of these being when *k*_+–_ > *k*_+–,*crit*_, when repression is fast enough to decrease the total number of available conjugators, which effectively slows down the adaptation process to the antibiotic and leads to a decrease in the effective growth rate. The second region is defined by the region of 0 < *k*_+–_ ≤ *k*_+–,*crit*_, when repression increases the effective growth rate by limiting HGT-driven mutation.

One of our main interests in this model was to evaluate possible fitness costs and fitness restoration, as in our previous static landscape model we have found that a bacterial population could regain fitness loss due to plasmid and/or resistance maintenance by modulating the repression of conjugation. Surprisingly, we found that in a dynamic landscape the effective growth rate of the population can be increased due to repression (Figure 7). However, since the effective growth rate isn’t mathematically formulated as the population mean fitness that we employed in our previous work, we suggest that this effect may have implications for the mean fitness as well. We quantified the difference between a population with no repression (although still able to conjugate) and a population that has reached the optimal value of repression as we have derived it for the case of *γρ* →∞. We note that this value should be identical for the limit case of *γρ* → ∞,*Κ_D_* →∞.

From an evolutionary perspective, this agrees with the idea that was previously suggested, namely that repression is a compensation mechanism for bacteria whose purpose is to reclaim fitness loss whenever a time comes when a current environmental stress (i.e., antibiotic) diminishes (Lenski 2010). Repression could also be applied as a strategy to mitigate the fitness cost involved in hosting a plasmid and/or antibiotic resistance and relaxed if the population encounters the same antibiotic in the future, serving as a “genetic memory” for an environmental response of the population as was previously suggested from experimental studies (Dionisio *et al.* 2005) and computational studies (Haft *et al.* 2009).

Our secondary derivation of our model by *γρ* (Section D) has allowed us to compute terms for the transition values of the de-repressed conjugators to repressed conjugators regions for two separate cases, when all conjugators (repressed and de-repressed) have adapted to an antibiotic and when all conjugators have not adapted but maintain a positive effective growth rate (detailed in Appendices C and D). However, only the latter of these cases is independent of the peak shift cycle time while the first case is not. Regardless, both terms are inversely related to the population of the repressed conjugators 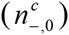 which we have detailed the full analytical solution in (Section D of the SI). We note that a similar phenomenon was observed in our previous models (Raz and Tannenbaum 2010; Atsmon-Raz and Tannenbaum 2014) as well as in other studies (Park and Deem 2007; Munoz *et al.* 2008).

To summarize, we have gained new qualitative insights into the significance of repression in the dynamics of antibiotic resistance through the construction and analysis of the model which we have discussed in this paper. We believe that the basic logic of this model could be applied to other biological systems which confer host/parasite dynamics with gained (or lost) fitness costs such as the Australian sheep blowfly *Lucilia cuprina* (Mckenzie *et al.* 1982), HIV drug resistance and evolution (Buckheit 2004) and evolution of herbicide resistance in plants(Vila-Aiub *et al.* 2009). More recently, studies have demonstrated that HGT is relevant in the proliferation of micrometastatic cells (Trejo-Becerril *et al.* 2012), giving further importance to this model regarding the study of cancer.

However, while theoretical models such as our own may provide us with more qualitative insights into the various processes of evolutionary dynamics, experimental characterization and validation of their parameters is essential to give a quantitative context to such models. Our current work demonstrates a steady development in the application and significance of quasi-species theory since Eigen’s original work (Eigen 1971) to the present day which we believe is a sign for the full potential of theoretical modelling within this framework for biological processes.

For future work, we will need to take into account several key biological features of conjugation, consideration and regulation of multiple plasmid copies in each bacterium within the population, plasmid compatibility classes, generation-dependent degradation of conjugation rates (Dahlberg and Chao 2003; Baltrus 2013), the effect of plasmid-bacterium coevolution on conjugation (Harrison and Brockhurst 2012), and most importantly non-single-point-mutation fitness landscapes. Finally, we aim to provide an in-depth analysis on the implications of horizontal gene transfer on the error-catastrophe phenomenon in a dynamic landscape.

## Acknowledgements

This paper was completed after the untimely death of Professor Emmanuel David Tannenbaum (of blessed memory) in May 2012, the author accepts all responsibility for the integrity of the data, analysis and results as published.

Emmanuel was an inspiring adviser, a great friend and an exceptional person in all aspects of his life. His kindness, brilliance, and humanity will always be remembered and treasured by those of us who had the honor of knowing and working with him.

